# Between a rock and a hard place: Effects of land-use change on rock-dwelling animals of lateritic plateaus in the northern Western Ghats

**DOI:** 10.1101/2023.01.12.523866

**Authors:** Vijayan Jithin, Manali Rane, Aparna Watve, Varad B. Giri, Rohit Naniwadekar

**Author notes:** Correspondence: Rohit Naniwadekar, Nature Conservation Foundation, Mysore, Karnataka, India.; Vijayan Jithin, Nature Conservation Foundation, Mysore, Karnataka, India.

## Abstract

Open natural ecosystems like lateritic plateaus, are undergoing rapid transformation with very poor understanding of these impacts on the threatened and endemic biodiversity. The unprotected, low-elevation lateritic plateaus of the northern Western Ghats are case to the point, as they have high endemicity but remain unprotected under Indian law. We aimed to understand the impact of the conversion of the natural lateritic plateaus to agroforestry and paddy cultivation on biodiversity. We compared the prevalence of two species of endemic herpetofauna of the northern Western Ghats (*Gegeneophis seschachari* and *Hemidactylus albofasciatus*) and a widespread snake (*Echis carinatus*) and the composition of other rock-dwelling animals across 12 undisturbed plateau sites and 10 sites each in agroforestry plantations and abandoned paddies on plateaus using time-constrained searches. We had 5738 encounters with 38 different animal species/groups. We found that the abundance of large rocks, which were the most-preferred size class of rocks by animals, was higher in abandoned paddy compared to plateaus and orchards. However, the prevalence of *H. albofasciatus* and *E. carinatus* was highest on undisturbed plateaus. Contrastingly, the prevalence of *G. seshachari* was significantly higher in abandoned paddy than undisturbed plateau or orchards. Non-metric multi-dimensional analysis showed that the assemblage of rock-dwelling fauna differed significantly across the three land-use types. Despite being adapted to persist in extremely variable climates on lateritic plateaus, we find that multiple species/groups are vulnerable to land-use change. However, *G. seshachari* and a few other taxa appear to benefit from certain kinds of land-use change, highlighting the context-specificity in species responses. While multiple studies have determined the impacts of forest conversion to other land-uses, this is one of the first studies to determine the impacts of the conversion of rocky outcrops, thereby highlighting the conservation value of habitats that are often classified as wastelands.

## INTRODUCTION

Open Natural Ecosystems, which include rock outcrops, sparsely treed desert ecosystems, savanna grasslands, and savanna woodlands, occupy more than a quarter of the world’s land area and are highly diverse (Dinerstein et al., 2017; Bond, 2019). While providing critical livelihood resources for humans, these highly threatened habitats are home to many endemic species (Singh et al., 2006; Bond, 2019). In India, open natural ecosystems that cover more than 10% of the country’s geographical area are unfortunately classified as ‘wastelands’ (Madhusudan & Vanak, 2022). These unrecognised and unprotected habitats are highly threatened due to land-use conversion (Madhusudan & Vanak, 2022). While most studies focus on land-use change impacts on forested habitats, our understanding of how land-use change affects biodiversity inhabiting the unique lateritic plateaus is very poor.

The low-elevation ferricretes or rock outcrops or lateritic plateaus of the northern Western Ghats, a type of open natural ecosystem with a significantly large part of the habitat as exposed rock, are being converted to other land-uses at an alarming rate (Bhattacharya et al., 2019). Lateritic plateaus in the northern Western Ghats harbour unique flora and fauna compared to wooded habitats in the neighbourhood, thereby forming habitat islands in the larger landscape (Porembski & Barthlott, 2000; Watve, 2013). Plateaus are critical habitats for multiple taxa, and new species of herbaceous plants, invertebrates, and vertebrates continue to be discovered in these habitats (Sayyed, Pyron, & Dileepkumar, 2018; Joshi, Karanth, & Edgecombe, 2020; Kulkarni, Shigwan, et al., 2022). More than 50% of plant species on the lateritic plateaus are endemic (Kulkarni, Shigwan, et al., 2022). These lateritic plateaus are mostly privately-owned or governed by the revenue department of the Government of India. Conversion to farms is one of the main threats to these lateritic plateaus. (Thorpe, 2018; Kulkarni, Shigwan, et al., 2022). Parts of these lateritic plateaus have been historically converted to paddy plantations and, more recently, into mango or cashew orchards (Bhattacharyya et al., 2019). The landscape is mosaic, with relatively less disturbed plateaus, forest patches, paddy, orchards, and human settlements, making a ‘constructed landscape’ in a socio-ecological sense. While land-use change in forested ecosystems has been known to impact animals in different ways including change in the community composition (Bobrowiec & Gribel, 2010), functional diversity (Etard, Pigot, & Newbold, 2022), species interactions (Tylianakis et al., 2008), facilitating invasions (Sánchez-Ortiz et al., 2020), risk from climate change (Williams & Newbold, 2020), the ecology of these rocky outcrops and the impacts of land-use change on rock outcrop biodiversity remains poorly studied (Kulkarni, Shigwan, et al., 2022).

Cold-blooded animals respond strongly to abiotic factors (Hofer, Bersier, & Borcard, 1999; Naniwadekar & Vasudevan, 2007) and land-use change (Gardner, Barlow, & Peres, 2007; Cordier et al., 2021). Seemingly homogeneous and exposed lateritic plateaus offer microhabitats like boulders that act as important refuges protecting animals from extreme weather and predation (Shah et al., 2004; Webb, Pringle, & Shine, 2009). Land-use change may significantly alter the quality and availability of these microhabitats. Large-scale movement of rocks impacts the availability of microhabitats for animals and, consequently, the abundance of animals (Webb & Shine, 2000; Goldingay & Newell, 2017). In the northern Western Ghats, especially in the coastal belt, large rocks sourced from the lateritic plateau are used in making compound walls, lining paddy fields and plant pits in orchards (Fig. 1; Appendix S8 & S9). In addition, soil is added to these plateaus for paddy cultivation altering the substrate. While previous studies have focused explicitly on boulder removal impacts on biodiversity (Shine et al., 1998; Pike et al., 2010), habitat conversion impacts on microhabitat availability and animal use are poorly understood. It is critical to determine the responses of cold-blooded animals, particularly the endemic and threatened species, to habitat conversion and altered microhabitat availability.

**Figure 1.**
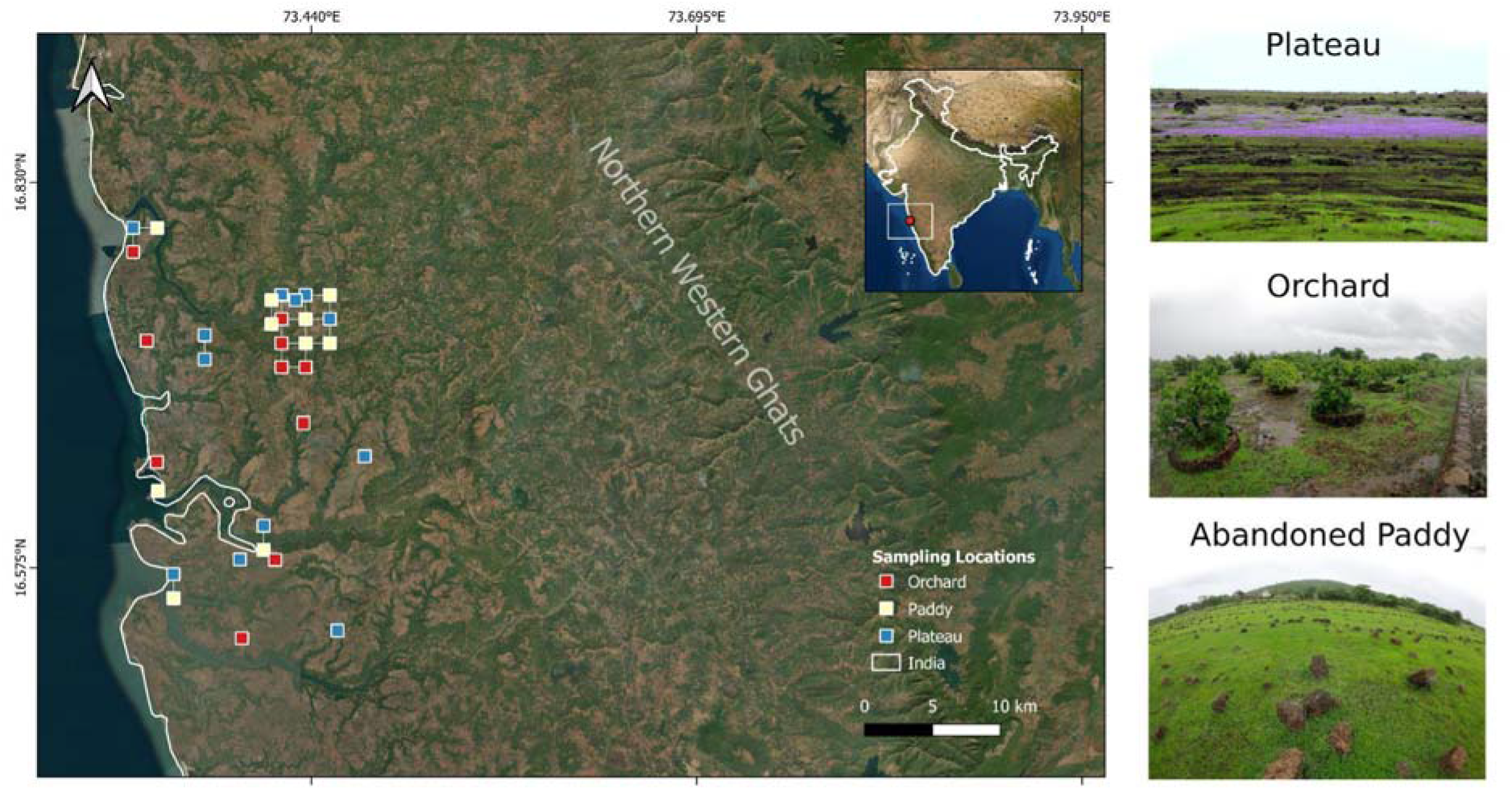
Map showing sampling sites across different land-use types on lateritic plateaus in the northern Western Ghats region of Ratnagiri District of Maharashtra in India. Satellite imagery source: Esri, Maxar, Earthstar Geographics, and the GIS User Community.

The boulders on low-elevation lateritic plateaus in the northern Western Ghats provide refuge to the ‘Vulnerable’ White-striped viper gecko *Hemidactylus albofasciatus*, ‘Data Deficient’ Seshachari’s caecilian *Gegeneophis seshachari*, Saw-scaled viper *Echis carinatus* among other taxa. *G. seshachari* is a fossorial caecilian endemic to the northern Western Ghats found in forested and plateau ecosystems. This lesser-known amphibian is encountered during the short-rainy season of the northern Western Ghats under rocks on the plateau, loose soil under boulders, and agricultural soil of approximately 10 cm depth (Gower, Giri, & Wilkinson, 2007; Katwate & Apte, 2019). It was re-discovered 36 years after the holotype collection from a mixed plantation habitat near a lateritic plateau (Gower, Giri, & Wilkinson, 2007). *H. albofasciatus* is an uncommon small, slender gecko found only in the open, rocky areas of lateritic plateaus of Maharashtra (Grandison & Soman, 1963; Amberkar, 2022). This ground-dwelling, nocturnal species is known to hide primarily under the rocks during the daytime and is patchily distributed (Gaikwad et al., 2009; Mirza & Sanap, 2012). The species is facing continuing decline due to rock collection for construction, stone quarrying, livestock grazing and mining (Srinivasulu & Srinivasulu, 2013). Saw-scaled viper, locally known as ‘Phoorsa’, occurs in forests, shrublands, grasslands, rocky areas and deserts (Ananjeva et al., 2021). However, the species can be found commonly under the rocks in plateaus during the monsoon (Sengupta et al., 1994). The species was heavily harvested for antivenin production, and the government organised destruction campaigns in the past, considering the health hazard to people (Sengupta et al., 1994; Whitaker & Whitaker, 2012). Between 1971 - 1991, almost 60,000 snakes were collected for anti-venin production near our study area (Sengupta et al., 1994). Apart from the three taxa, several invertebrates, including spiders, scorpions, centipedes, beetles, and ants, are also commonly found under these rocks.

In this study, we compared, across less-disturbed lateritic plateaus (reference sites), orchards and abandoned paddy, 1) the availability of different-sized rocks, 2) the occurrence of *G. seshachari, H. albofasciatus*, and *E. carinatus*, and 3) composition and abundance of other vertebrate and invertebrate taxa. Given that large rocks are used extensively in modified habitats, we expected that modifying the original plateau habitat to paddy and orchards would impact the abundance of rocks of the more preferred size classes (larger rocks) across land uses. In addition, land-use change will alter the occurrence of the caecilian, gecko and viper due to altered microhabitats. We expected differences in the composition and abundance of rock-dwelling faunal communities across the three land-use types.

## MATERIALS AND METHODS

### Study Area

The Western Ghats is among the eight ‘hottest’ biodiversity hotspots due to high endemism (particularly of herpetofauna) and the intensity of anthropogenic threats (Myers et al., 2000). Vast expanses of ferricretes and basalt plateaus characterise the northern portions of the Western Ghats. The low-elevation ferricretes (25-200 m asl) occur from the sea coast to the foothills, and the high-level ferricretes (800-1400 m asl) occur on high-level laterites to the east of the crestline of Western Ghats (Watve, 2013). Such vast lateritic plateaus are absent in Western Ghats’ central and southern portions (Kulkarni, Shigwan, et al., 2022). Composition of amphibians and aquatic vegetation differs between the high and low-elevation plateaus (Thorpe et al., 2018; Kulkarni, Vijayan, et al., 2022). The northern portion of the Western Ghats is more seasonal than the southern part, with rains primarily restricted to the four months of monsoon from June to September (Watve, 2008, 2013). Heavy rains in the monsoon transform the dry and exposed habitat into a lush green carpet of herbaceous vegetation. Hot and dry conditions during the dry season and water-logging during the monsoon expose animals inhabiting the rocky plateaus to extreme environments (Watve, 2008). The loose rocks on the lateritic plateaus provide refuge to a diverse array of invertebrate and vertebrate fauna (Thorpe & Watve, 2015).

We conducted the study in the low-level lateritic plateaus of Ratnagiri, in the Konkan region of Maharashtra state in west India (16°31’-16°48’N; 73°19-73°29’E; Fig. 1). These plateaus are locally known as ‘*sada*’ or ‘*jambha kaatal*’. The elevation of the sampled plateaus ranged between 24-197 m asl. The Arabian sea and the Western Ghats escarpment are to the west and east of the plateaus, respectively. The sampling locations on lateritic plateaus were privately-owned areas. Portions of the plateaus have been modified to paddy fields and cashew/mango orchards. Traditionally, paddy is either grown in existing depressions on plateaus that have accumulated soil or by dumping soil from neighbouring areas on the plateau. Large rocks from the plateaus are lined along the paddy fields to prevent soil run-off. Many paddy fields have been abandoned, given human migration to urban centres driven by stagnation in agriculture, poor returns from farming and other socio-economic reasons (Yamin, 1989). However, the abandoned paddy fields continue to harbour a layer of soil and loose rocks in high numbers (Fig. 1). In mango/cashew orchards on plateaus, the rocks are blasted with explosives, and the pits are filled with soil and lined with rocks (Bhattacharyya et al., 2019). Mango/cashew saplings are planted in the pits. Mangos grown on plateaus fruit at a desired time, well before the onset of monsoon, and are supposedly sweeter, fetching a higher price (Bhattacharyya et al., 2019). This has resulted in the rapid expansion of mango orchards, with more than 25,000 ha of lateritic plateaus now under mango orchards (Bhattacharyya et al., 2019). We sampled such abandoned paddy fields and orchards along with existing natural lateritic plateaus. The relatively less-disturbed plateaus were our reference sites. The orchards included mango and mixed mango-cashew orchards. The paddy fields sampled in the study were abandoned for at least two years and had a soil depth of at least 10 cm.

### Sampling

Between June to September 2022, coinciding with the monsoon period, we conducted one-hour time-constrained searches at 32 sites spread across 11 plateaus and three land-use types (12 natural lateritic plateaus (reference sites), 10 abandoned paddy fields and 10 orchards) (Fig. 1; Appendix S1). Time-constrained search enables the detection of more species and individuals per site compared to area-controlled plot methods (Kadlec, Tropek, & Konvicka, 2012). Given that our target species are rarely encountered, and since we were interested in understanding the micro-habitat use, we felt that time-constrained searches were a better strategy than belt transects. One observer (VJ) conducted the searches between 0900 - 1700 hr during the daytime, well after sunrise and before sunset. The sampling effort was similar across the three land-use types (10 hours each in orchards and abandoned paddy and 12 hours on plateaus) (Appendix S1). We walked from a starting point away from the road or edge of the habitat turning all the rocks encountered in the 1 hr walk. The general direction of the walk ensured that we remained within the same land-use type even after one hour. For each turned rock, the observer recorded the size class of the rock. We classified the rocks into small (< 5 cm in diameter of the planar surface), medium (5-10 cm) and large (> 10 cm) sizes based on the observation that mostly rocks of >10cm size were moved from plateaus for various purposes. All large rocks that we could manually move were checked. We immediately placed all the rocks in their original positions after recording the faunal community. The observer recorded all visible fauna, including *G. seshachari*, *H. albofasciatus*, and *E. carinatus*.

Apart from the three focal animals, all individual animals were counted, except in the case of ants (Formicidae), termites (Termitoidae) and mites (Acariformes) as they occurred in very high numbers making it difficult to count the exact number of individuals. Species-level identification in the field was difficult for *Minervarya* spp. froglets. Among arthropods, there was confusion in identifying one insect species that belonged to Dermaptera but was misclassified as Orthoptera. Arthropods were identified to the order or family level wherever possible.

### Analyses

We performed all analyses in R (v. 4.2.2) (R Core Team, 2022). We used a generalised linear model with a negative-binomial error structure (since Poisson error structure indicated overdispersion) to determine the differences in the number of rocks encountered across the different land-use types (lateritic plateaus, abandoned paddy and orchards). Since rocks, particularly large- and medium-sized ones, are moved to paddy from lateritic plateaus, we expected interactive effects between rock size and land-use type. Predictors whose 95% CI on the estimated coefficients did not overlap zero were thought to significantly influence the response variable. We estimated the marginal means and assessed pairwise contrasts for rock and land-use types in the model with Tukey’s method for *p*-value adjustment. We estimated Nagelkerke’s pseudo *R*^2^. We used the R packages ‘MASS’, ‘emmeans’, and ‘performance’ for this analysis (Venables & Ripley, 2002; Lüdecke et al., 2021).

We used generalised linear model with binomial error structure and each time-constrained search as sampling unit to determine if the occurrence of the three focal herpetofaunal species (*G. seshachari*, *H. albofasciatus* and *E. carinatus*) differed across the three land-use types. The analysis was carried out separately for each species. For *G. seshachari*, which was detected across the three land-use types, we used the regular binomial GLM in ‘MASS’ package. For *H. albofasciatus* and *E. carinatus*, which were never detected in abandoned paddy, we initially assessed separation and infinite maximum likelihood estimates for log-binomial regression using the package ‘detectseparation’ (Kosmidis, Schumacher, & Schwendinger, 2022). Since we detected separation and biased estimates, we used package ‘brglm2’ (Kosmidis & Firth, 2021) for bias reduction with a mixed bias-reduction adjusted scores approach to model the influence of land-use type on these two species. To see if the focal species preferred large-sized rocks over medium-sized ones, we used the Chi-squared test for independence.

We visualized the co-occurrence of all taxa across the land-use types, and rock-size classes using the package ‘UpSetR’ (Gehlenborg, 2019). To determine if the composition of the rock-dwelling community of lateritic plateaus differed across land use types, we used the *metaMDS* function of ‘vegan’ package (Oksanen et al., 2022) with the presence-absence data (since we did not have abundance information for ants and mites) of the taxa recorded across sites. Since the two-dimensional stress value was higher than 0.2, we used a three-dimensional ordination with the Bray-Curtis dissimilarity index. We validated the assumption of homogeneity of multivariate dispersion using *betadisper* function and tested for the differences between the land-use types using the Permutational multivariate analysis of variance (PERMANOVA) using the *adonis* function in package ‘vegan’. To assess the influence of land-use change on taxa with more than 100 detections, we used generalised linear models with negative-binomial error structure. We assessed pairwise contrasts for all three land-use types in each model as described before.

## RESULTS

### Rock availability across land-use types

We turned 7179 rocks across the different land-use types in 32 hr (and sites) across 11 different lateritic plateaus. We found that the model that examined variation in rocks of different size classes across the different land-use types explained significant variation in the observed data (Nagelkerke’s Psuedo *R*^2^ = 0.85). The interaction term between rock size and land-use type was significant (Fig. 2a and b; Appendix S3a). Compared to the less-disturbed lateritic plateaus, in the abandoned paddy, there were fewer small rocks but higher numbers of large rocks (Fig. 2a and b and Appendix S3b). The numbers of small-, medium- and large-sized rocks did not differ between orchards and less-disturbed lateritic plateaus (Fig. 2a and b; Appendix S3b).

**Figure 2.**
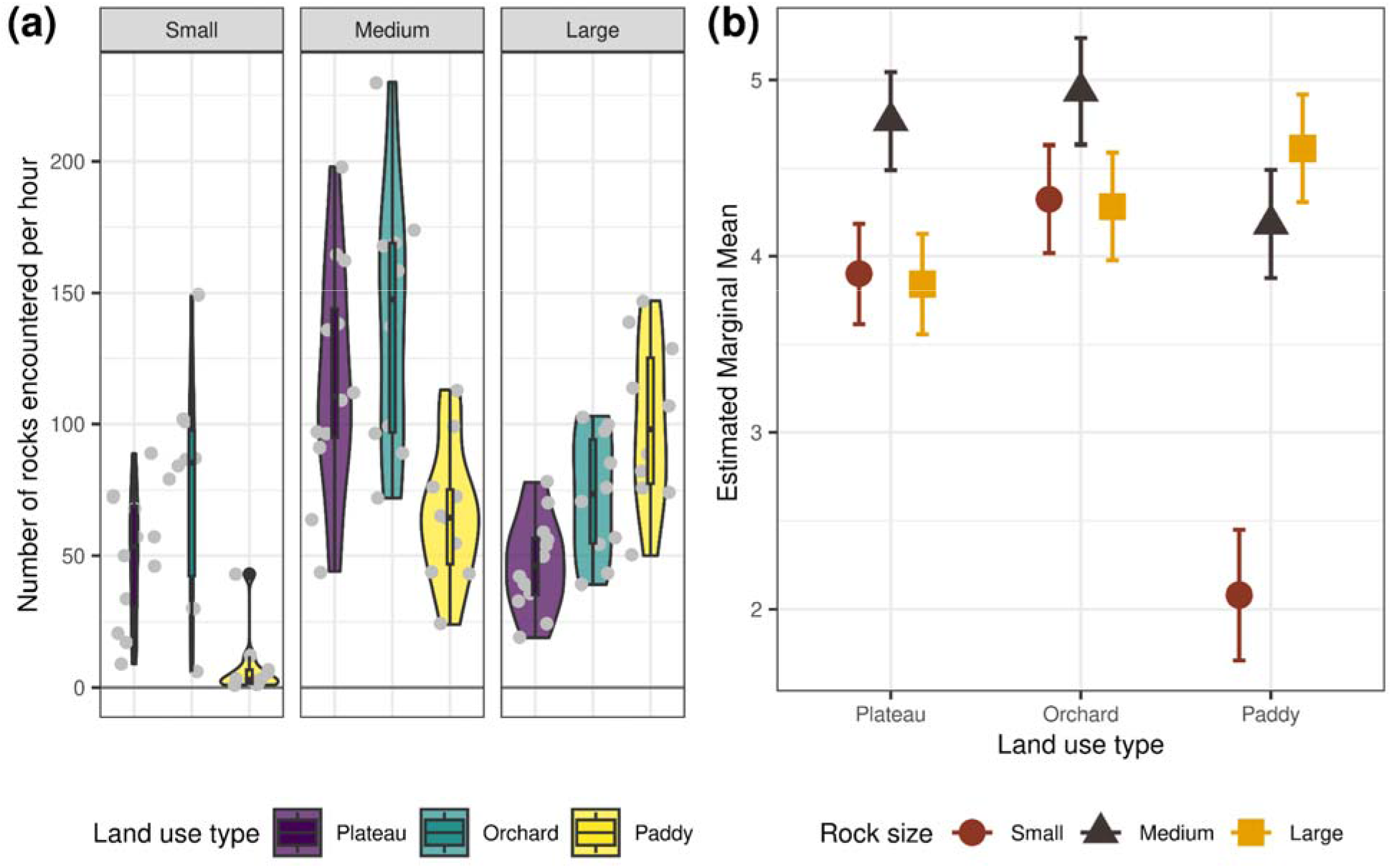
(a) The violin plot shows the rock encounter rates (per hour) of small- (< 5 cm diameter), medium- (5-10 cm diameter) and large-sized rocks (> 10 cm) across the three land-use types. The grey dots are individual data points. While the number of small- and medium-sized rocks was least in paddy, the number of large-sized rocks was highest in paddy; (b) Estimated marginal means with 95% CI from the generalized linear model used to assess the influence of land-use type on the encounter rate of rocks.

### Prevalence of caecilian, gecko and viper across land-use types

We had 35 detections of *G. seshachari* from three lateritic plateaus, 24 detections of *H. albofasciatus* from seven plateaus and 22 detections of *E. carinatus* from nine plateaus. We never detected these three species under small-sized rocks. The probability of occurrence of the caecilian was significantly higher in abandoned paddy than in less-disturbed plateaus (Fig. 3; Appendix S4). However, we never detected the gecko and viper in paddy (Fig. 3A). The probability of occurrence of *H. albofasciatus* and *E. carinatus* was significantly higher in less-disturbed plateaus compared to the orchards and abandoned paddy (Fig. 3; Appendix S4). All three animals used large-sized rocks more than medium-sized ones (*G. Seshachari*: □^2^ = 12.6, *df* = 1, *p* < 0.001; *H. albofasciatus*: □^2^ = 6, *df* = 1, *p* = 0.01; *E. carinatus*: □^2^ = 6.55, *df* = 1, *p* = 0.01; Fig. 3A).

**Figure 3.**
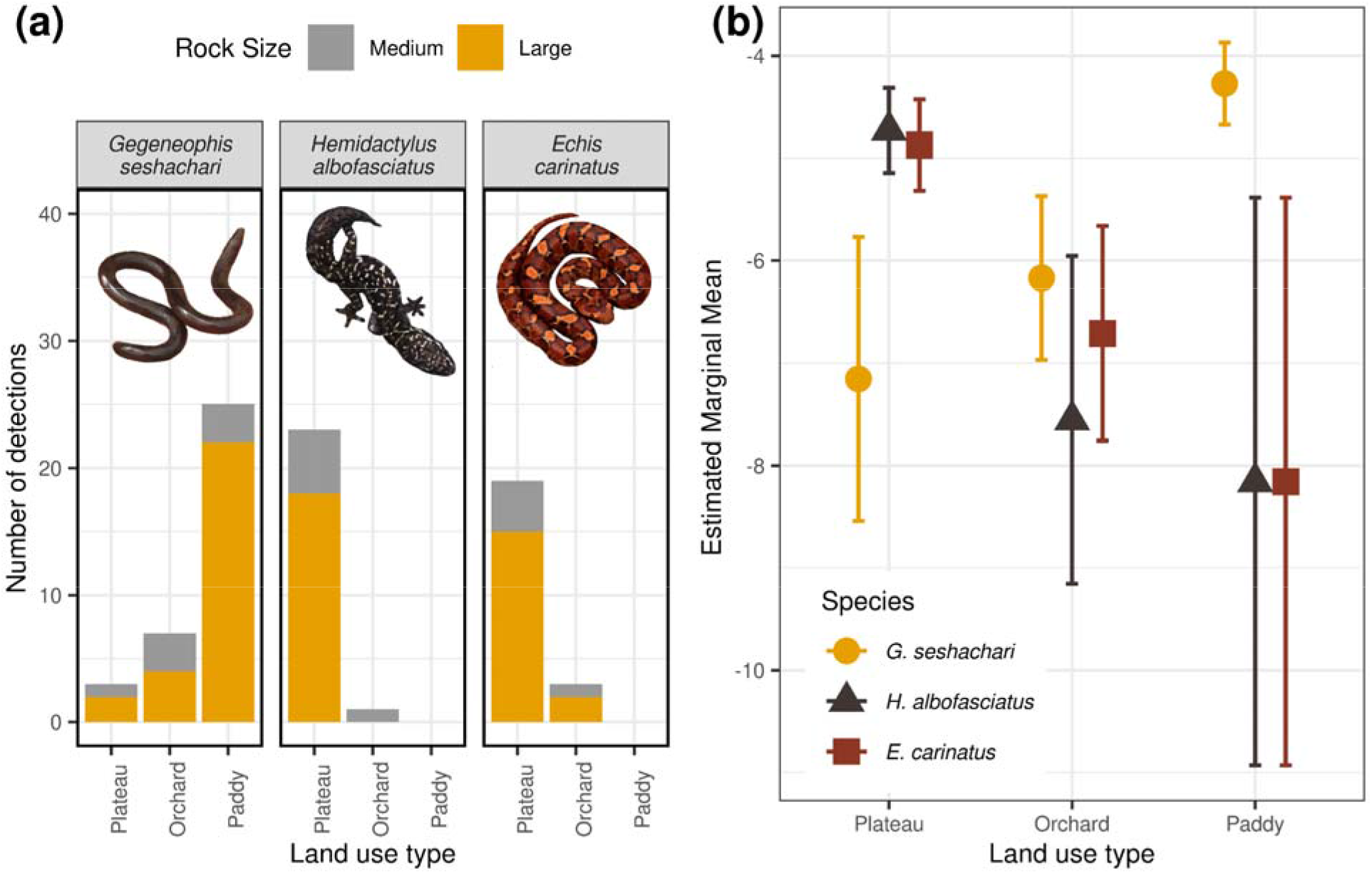
(a) Number of detections of *Gegeneophis seshachari, Hemidactylus albofasciatus*, and *Echis carinatus* across land use types and rock size classes; (b) Estimated marginal means with 95% CI from the generalized linear models used to assess the influence of land-use type on the proportion of detection of focal species. Photographs by V. Jithin and A. Gadkari.

### Composition of the animal community under rocks across land-use types

Animals were detected under almost 45% of the rocks turned (58%, 44% and 24% under the large, medium and small rocks, respectively). Apart from the *G. seshachari*, *H. albofasciatus*, and *E. carinatus*, we recorded eight genera of frogs, three genera of lizards, two species of snakes and one mammal. We documented 38 taxonomic groups spanning 10 classes and more than 20 orders under rocks across different land uses from 5738 encounters (Appendix S2 and S5). The most numerically dominant groups of animals that we found under the rocks were Insecta (2937), followed by Clitellata (861), Arachnida (671), Malacostraca (471), Chilopoda (320), Amphibia (283) and Gastropoda (106). Darkling beetles (*Gonocephalum* sp.), earthworms (Oligochaeta), ants, crickets and earwigs were among the most abundant animals. The co-occurence plots showed that paddy harboured six exclusive taxa, whereas less-disturbed plateaus had two, and orchards had none. Nine taxa occurred across all land-use types and rock-size classes (Appendix S6). NMDS analysis demonstrated that the animal communities found under rocks differ significantly across land-use types (PERMANOVA *R*^2^ = 0.21, df = 2, *F* = 3.82, *p* = 0.001; Fig. 4). Animal composition under rocks in paddy was very distinct from lateritic plateaus (Fig. 4).

**Figure 4.**
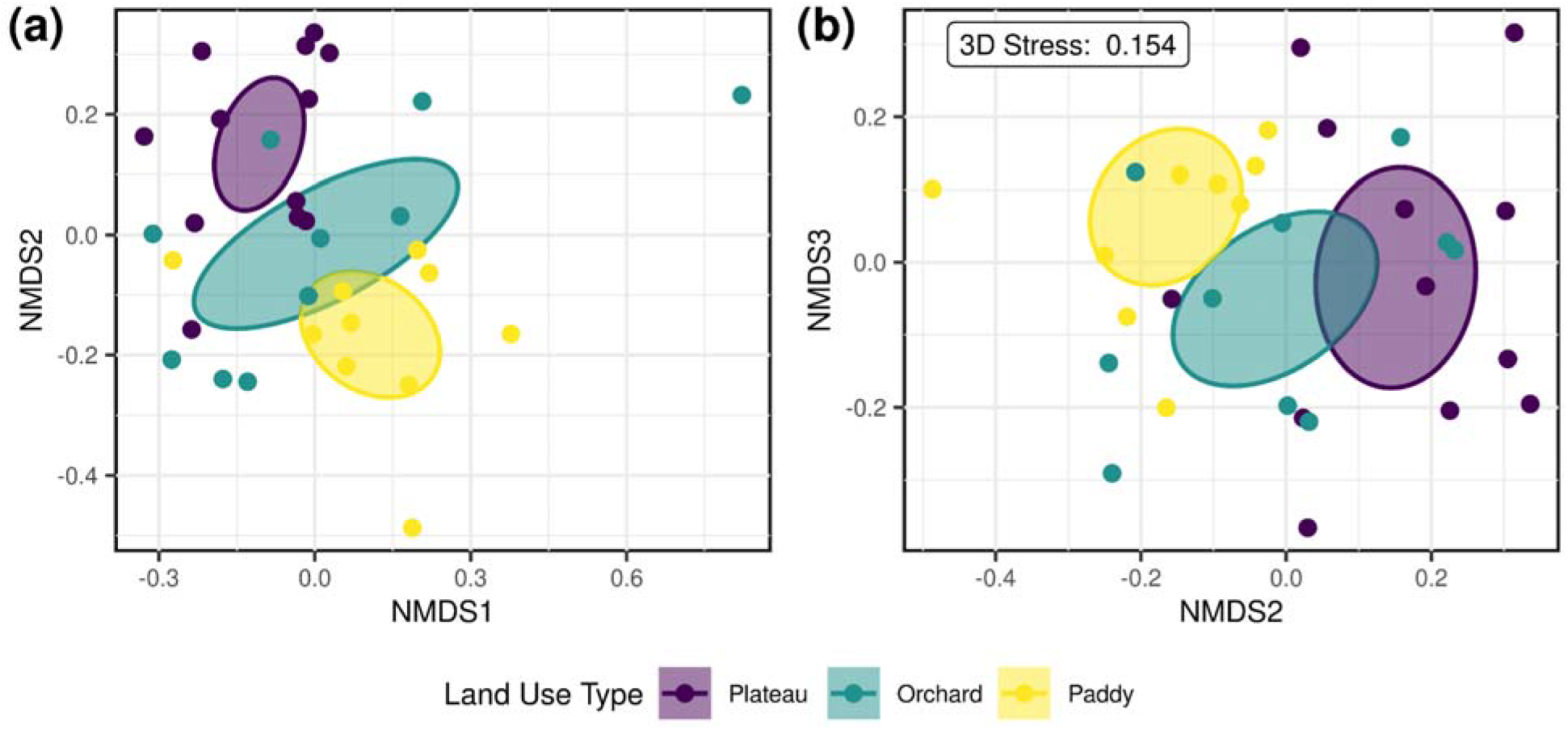
Non-metric-multidimensional-scaling in three dimensions showing dissimilarities between the three land-use types (less-disturbed plateaus, orchards, abandoned paddy) in the overall taxonomic composition of rock-dwelling animals in the lateritic plateaus of the northern Western Ghats. Figure (a) represents the first two axes (NMDS1 & NMDS2) and (b) represents the second and third (NMDS2 & NMDS3) axes. The ellipses indicate multivariate 95% confidence intervals around the group centroid.

Less-disturbed plateaus had a significantly higher abundance of Araneae and *Gonocephalum* sp. than orchards; and Araneae, *Hottentotta* sp. and Dermaptera-Orthoptera groups than abandoned paddy (Fig. 5; Appendix S7). The abandoned paddy had a significantly higher abundance of Isopoda and Blattoidae than orchards; and Blattoidae than less-disturbed plateaus (Fig. 5; Appendix S7). The orchards showed a significantly higher abundance of *Hottentotta* sp. and Araneae than the abandoned paddy (Fig. 5; Appendix S7). The abundance of *Minervarya* spp., Oligochaeta, Scolopendridae, Formicidae, Gastropoda, and Scarabaeidae groups did not show any significant difference between any land-use types (Fig. 5; Appendix S7).

**Figure 5.**
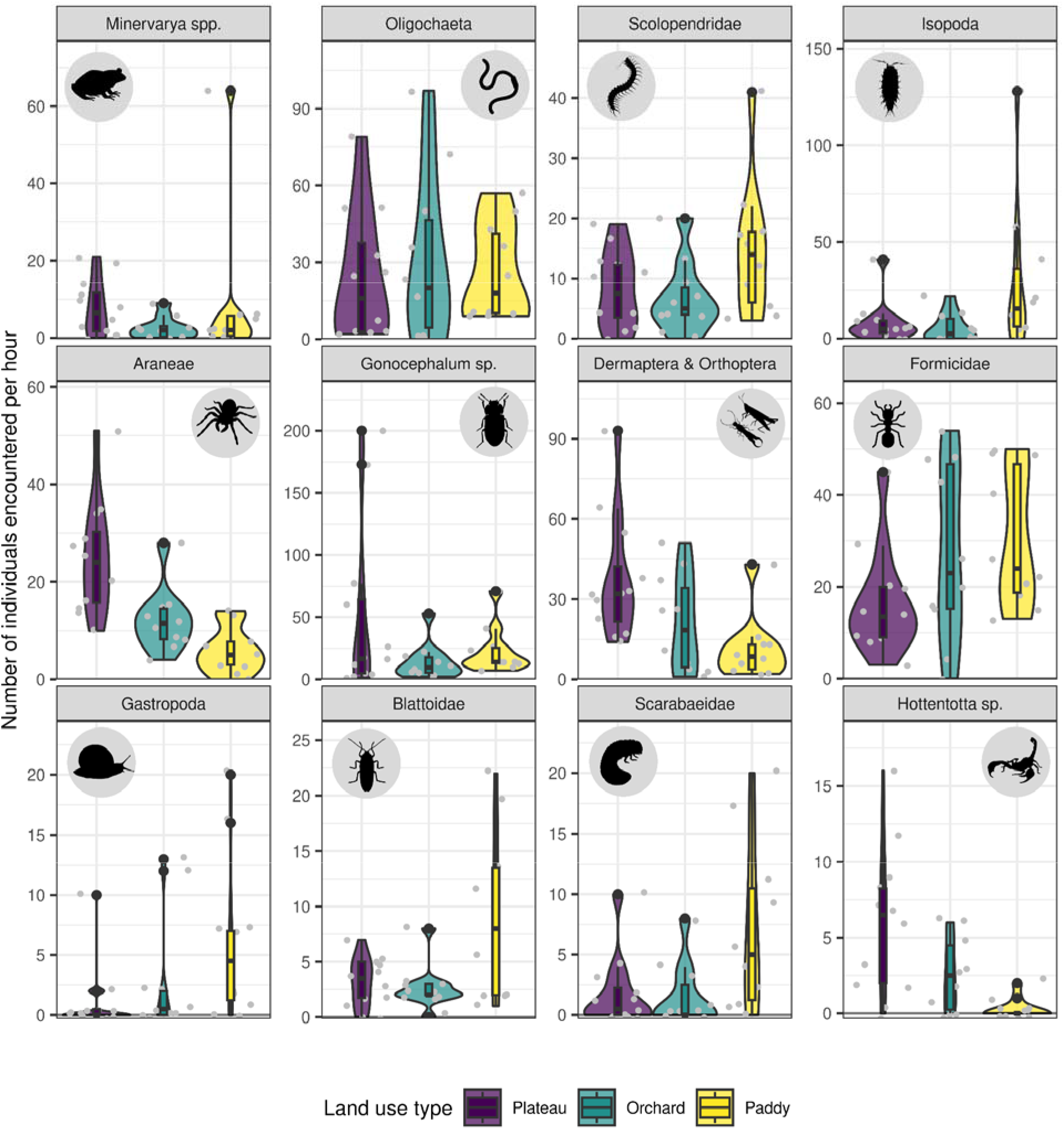
Violin plots showing the encounter rates (per hour) of animals with greater than 100 detections during the survey across the three land-use types. The grey dots are individual data points. Note that the numbers on the y-axis for Formicidae are the number of group encounters (per hour) (see the text for details).

## DISCUSSION

Land-use change is among the biggest drivers of herpetofaunal declines (Ficetola et al., 2015; Doherty et al., 2020). Land-use change reduces the quantum of suitable habitat, thereby negatively impacting the diversity and abundance of amphibians and reptiles, especially specialist species (Ficetola et al., 2015). However, previous studies have mainly examined the impacts of the conversion of forested landscapes to other land-uses (Kurz et al., 2014). We demonstrate the impacts of land-use conversion of the unique lateritic plateaus on select threatened and endemic species and on the community composition of rock-dwelling fauna. Land use change of lateritic plateaus to paddy resulted in a high influx of larger rocks in abandoned paddy. The two endemic species exhibited contrasting responses to land-use change. The conversion of lateritic plateaus to agroforestry plantations and paddy negatively impacted the threatened and endemic *H. albofasciatus* and the generalist *E. carinatus*. Interestingly, *G. seshachari*, an endemic amphibian that occurs in forests and plateaus, was more prevalent in the abandoned paddy than in less-disturbed plateaus and orchards. Habitat conversion significantly changes the composition of rock-dwelling faunal communities on the lateritic plateau. This is one of the first studies to examine the impacts of land-use change on rock-dwelling fauna across a wide taxonomic breadth. The study also generates valuable baseline information on endemic and threatened species inhabiting the unique lateritic plateau habitat of the Western Ghats Biodiversity hotspot, one of which (*G. seshachari*) has been identified as ‘Data Deficient’ by the IUCN.

### Large rocks as a refuge

Large-sized rocks on outcrops, which experience smaller daily temperature amplitudes, and maintain temperatures for a longer duration, may be favoured by cold-blooded animals (Goldsbrough, Hochuli, & Shine, 2003). In the rock outcrops of Australia, broad-headed snake *Hoplocephalus bungaroides* and its principal prey, the Velvet gecko *Amalosia lesueurii*, depend on rocks with distinct thermal profiles as shelter sites (Webb & Shine, 1998). We found that the caecilian, gecko and viper consistently preferred large rocks. Invertebrates were frequently encountered under large- and medium-sized rocks but were relatively rarer under small rocks (Appendix S5), highlighting the value of large rocks as important refuges on the open lateritic plateaus.

Just the presence of large rocks need not necessarily influence the occurrence of different animals. While the caecilian, gecko, viper and many other faunal groups preferred the large rocks, they still demonstrated contrasting responses to conversion to paddy. *E. carinatus* and *H. albofasciatus* prefer large rocks, yet they were not detected in paddy fields with the highest numbers of large rocks among the three land-use types. The substratum under rocks is also modified from hard rock to muddy soil with modification to paddy. Webb & Whiting, (2006) found that the probability of occurrence of snakes was higher under boulders on rock substrate than on soil substrate. The modified microhabitat and microclimate under large rocks can be unsuitable to rock-dwelling fauna, particularly the *E. carinatus* and *H. albofasciatus*, which may prefer the rocky substratum over muddy soils. Land-use affects both availability and the quality of microhabitats, an aspect which needs to be studied further.

Apart from lining the paddy, large rocks are also used extensively to construct compound walls and mark land borders among other activities (Mirza & Sanap, 2012; Thorpe & Watve, 2015). All of these activities reduce the availability of large rocks on neighbouring plateaus. This might be detrimental to the prevalence of rock-dwelling fauna, especially the viper and the gecko. Rock addition experiments could be carried out on lateritic plateaus, particularly in proximity to paddy fields, to determine species recovery. A socio-ecological study can be carried out to understand the zone of influence around the paddy fields from where large rocks are sourced by locals and its impacts on herpetofauna, which can provide valuable insights for restoration efforts in the future.

### Contrasting responses of reptiles and caecilian

Herpetofaunal communities respond differentially to habitat conversion to monocultures and polyspecific plantations (Trimble & van Aarde, 2014; Iglesias Carrasco, Medina, & Ord, 2022). Reptiles are more resilient to land-use change than amphibians (Fulgence et al., 2022). Most of the past research has focused on anurans with little information on the impacts of land-use change on caecilians (Valdez, Gould, & Garnham, 2021). The present study is one of the first to determine the responses of caecilians to land-use change. While the viper and the gecko are negatively affected by land-use change, the muddy soils of the abandoned paddy may have provided increased opportunity for burrowing and a more amenable cooler microclimate that may reduce water loss of the endemic and fossorial caecilian.

Generally, land-use change negatively impacts endemic and habitat specialist species while benefiting generalist species (McKinney & Lockwood, 1999; Newbold et al., 2018). Instances of endemics benefitting from land-use change, like the caecilian in our case, are not common. The endemic and forest specialist saproxylic beetles in the Santa Maria Islands continued to persist in plantations (Meijer, Whittaker, & Borges, 2011). Unlike the habitat specialist *H. albofasciatus*, *G. seshachari* is a habitat generalist found in forests and plateaus. The soil substrate under large rocks in abandoned paddy likely offers the fossorial amphibian a suitable micro-environment to bury. However, we must be mindful that only certain kinds of land-use change (like abandoned paddy in this case) likely provided a more favourable habitat for the caecilian. Farmland abandonment has varying consequences for biodiversity (Queiroz et al., 2014) as in this study. Also, given that the paddy fields have been abandoned, one question is whether we move the large rocks from abandoned paddy to plateaus. Further thought and research are required as moving rocks from abandoned paddy to lateritic plateaus might benefit the gecko and viper but may come at a cost for the caecilian.

### Other factors influencing reptile prevalence

In our study, the availability of large rocks was similar between plateaus and orchards, but gecko and viper abundance was lower in orchards. While the number of rocks did not differ significantly between orchards and plateaus, orchards in the lateritic plateaus change the microhabitat structure due to addition of trees in an otherwise treeless landscape (Amberkar, 2022). Disturbances in the habitat or alterations of the habitat structure can lead to differences in the abundance between natural and modified habitats (Gorissen, Greenlees, & Shine, 2017). Apart from the modified microhabitat, human persecution could also be responsible for the rarity of the viper. Humans fear venomous vipers and are persecuted when seen (Balakrishnan, 2010). *Echis carinatus* is one of the big four venomous snakes, which are responsible for almost 11,000 human deaths annually in India (Das, 2002; Mohapatra et al., 2011). While the lateritic plateaus are used as grazing grounds for cattle, humans are more likely to persecute vipers in modified habitats, given higher human activity. Vipers are supposed to be very common in the study area. This was one of the preferred regions for collecting vipers for anti-venin production (Sengupta et al., 1994). Two decades ago, a team of five members collected approximately 150 vipers in five hours from lateritic plateaus in the vicinity of our sampling area (pers. comm. Sanjay Thakur). However, in a comparable effort, we found only 22 vipers. This underlines the need to systematically evaluate how viper collection may have impacted their regional populations.

The most common prey item of *E. carinatus* and *Hemidactylus* sp. are arthropods (Cyriac & Umesh, 2021; Ghezellou et al., 2021); and of *Gegeneophis* are termites and earthworms (Measey et al., 2004). While land-use change did not alter the overall abundance of arthropods, we found a change in the community composition of rock-dwelling invertebrates. Replacement of Araneae, Dermaptera and Orthoptera with Scolopendrid centipedes, Oligochaetes, Formicids and Isopods likely resulted in this turnover in the communities. We had expected encounters of invertebrates to be lower in orchards compared to the less-disturbed plateau due to the heavy use of pesticides in orchards. But, except for the lower numbers of spiders and darkling beetles, we did not find any difference. However, further studies are required on the pesticide effect on invertebrates considering the seasonality in pesticide use. Future studies should aim to generate fine-scale information on gecko, caecilian, and viper diets through metabarcoding and compare the abundance of preferred prey items across land-use types in long-term to determine the potential role of prey availability in distributions of the gecko, viper and caecilian.

Information from locals suggested potential threats of illegal pet trade to the gecko, an aspect that needs to be studied in greater detail. Illegal pet trade, even of relatively newly described species, is one of the most pervasive threats to select species of geckos, including other species of viper geckos (Altherr & Lameter, 2020). Similarly, altered predation pressure associated with land-use change and its cascading effects on animals (Schwab et al., 2021) can be an interesting aspect to look at in the future. Given the harvest, persecution and other factors discussed above, it is critical to monitor populations of the vipers, gecko and caecilian, particularly in sites where they are relatively common. In this study, using a non-invasive method, we have established baselines of the rock-dwelling faunal community that can be useful for periodic monitoring.

### Conservation of lateritic plateaus

The low-elevation lateritic plateaus are not protected, have received poor conservation attention and are being rapidly converted to orchards. Proposed development projects like refineries and power plants have also been planned. Additionally, plateau rocks are being used extensively for multiple human activities. Therefore, there is a need for systematic biodiversity inventories of the lateritic plateaus, identifying key plateaus and working with local communities towards their conservation. Plateaus, where large rocks have been heavily depleted, habitat restoration through the introduction of aptly designed artificial rocks can be explored (Croak et al., 2010; Croak, Webb, & Shine, 2013). The information collected in this study, in conjunction with other socio-ecological information collected as part of other ongoing projects, will contribute to declaring select plateaus as ‘Biodiversity Heritage Sites’ with the due consent of local communities.

### Limitations of the study

Invertebrate taxonomy in the region is still not resolved as new species continue to be described. This did not allow us to identify different invertebrates at the species level. However, even with higher levels of taxonomic information, NMDS demonstrated significant compositional differences. Fine-scale taxonomic information will most likely exacerbate these differences. Since local communities extensively use all the plateaus, we did not have a completely undisturbed laterite plateau as a benchmark. Therefore, our inferences of different landscapes are, at best conservative, since undisturbed sites may have had much higher abundances of some of these animals.

## CONCLUSIONS

We demonstrate that land-use change in open natural ecosystems can have drastic impacts even on species adapted to persist in extreme and variable environments. Land-use change, while impacting specialist and generalist species, may also be advantageous to certain endemic species under certain circumstances. Land-use change is often seen in the light of the benefit to people due to agricultural intensification and development, though it results in biodiversity decline and the associated loss in ecosystem function and services to humans (Hooper et al., 2012; Newbold et al., 2015). Future studies need to systematically document the impacts of land-use change on biodiversity and ecosystem function in open natural ecosystems and identify thresholds that may facilitate the coexistence of humans and biodiversity.

## ACKNOWLEDGEMENTS

We thank Maharashtra Forest Department, particularly the Chief Wildlife Warden, Sunil Limaye, for giving us the necessary permits (Letter No. Desk-22(8)/WL/Research/CR-53(20-21)/3361/22-22) to conduct the study. We thank On the Edge (UK), The Bombay Environmental Action Group and The Habitat Trust (India) for funding this work. We are grateful to Poornima Viswanathan for helping identify arthropods. We thank Aditya Gadkari, Ajay Nachinekar, Chandrakanth Gurav, Harshad Tulpule, Kamalakar Gurav, Pooja Ghate, Poorva Joshi, Pradeep Dingankar, Rakesh Patil, Rashmi Karandikar, Ravindra Karandikar, Santosh Padhye, Shailesh Joshi, Suhas Gurjar, Sujan Dandekar, and Yash Vichare for support during fieldwork. RN thanks Anand Osuri, Kulbhushansingh Suryawanshi, Suri Venkatachalam, Madhura Niphadkar and Vijay Karthick for useful discussions.

## AUTHORS’ CONTRIBUTIONS

*VJ* and *RN* conceived the ideas and designed the methodology with inputs from AW and VG; *VJ* collected the data; *VJ* and *RN* analysed the data; *VJ and RN* led the writing of the manuscript with inputs from VG, AW and MR. All authors contributed critically to the drafts and gave final approval for publication.

## DATA AVAILABILITY STATEMENT

Data and codes used in this study will be uploaded on DataDryad on acceptance.

