## Supporting Files for "Between a rock and a hard place: Effects of land-use change on rock-dwelling animals of lateritic plateaus in the northern Western Ghats"

*Correspondence:

**Appendix S1.** Sampling effort across different plateaus and land use types

| **Land Use** | **Plateau Name** | **No. of Rocks Turned** | **No. of 1 hr Time-constrained Searches** |
| --- | --- | --- | --- |
| Plateau | Bakale | 150 | 1 |
|  | Barsu | 260 | 1 |
|  | Devache Gothane | 249 | 1 |
|  | Devi Hasol | 712 | 4 |
|  | Gaokhadi | 240 | 1 |
|  | Nanar | 305 | 1 |
|  | Niveli | 231 | 1 |
|  | Rundhye | 417 | 2 |
| Orchard | Ambolgad | 339 | 1 |
|  | Devache Gothane | 351 | 1 |
|  | Devi Hasol | 961 | 4 |
|  | Gaokhadi | 344 | 1 |
|  | Nanar | 299 | 1 |
|  | Niveli | 310 | 1 |
|  | Vetye | 268 | 1 |
| Paddy | Bakale | 161 | 1 |
|  | Devache Gothane | 190 | 1 |
|  | Devi Hasol | 936 | 6 |
|  | Gaokhadi | 270 | 1 |
|  | Musa Kazi | 186 | 1 |
| **Total** | | **7179** | **32** |

**Appendix S2.** Complete list of animals detected under the rocks turned across the land-use types with number of detections.

| **Class** | **Taxa** | **Number of detections** | | | |
| --- | --- | --- | --- | --- | --- |
|  |  | **Plateau** | **Orchard** | **Paddy** | **Total** |
| Amphibia | *Duttaphrynus melanostictus* | 1 | 1 | 0 | 2 |
|  | *Euphlyctis jaladhara* | 0 | 0 | 15 | 15 |
|  | *Gegeneophis seshachari* | 3 | 7 | 25 | 35 |
|  | *Hoplobatrachus tigerinus* | 8 | 0 | 0 | 8 |
|  | *Microhyla* cf. *nilphamariensis* | 2 | 0 | 2 | 4 |
|  | *Minervarya* spp. | 95 | 26 | 87 | 208 |
|  | *Polypedates maculatus* | 0 | 0 | 1 | 1 |
|  | *Sphaerotheca dobsonii* | 4 | 4 | 0 | 8 |
|  | *Uperodon mormoratus* | 0 | 1 | 1 | 2 |
| Arachnida | Acariformes | 0 | 0 | 4 | 4 |
|  | Araneae | 299 | 122 | 59 | 480 |
|  | *Sahyadrimetrus* sp. | 28 | 10 | 48 | 86 |
|  | *Hottentotta* sp. | 72 | 26 | 3 | 101 |
| Chilopoda | Scolopendridae | 98 | 68 | 147 | 313 |
|  | Scutigeromorpha | 3 | 0 | 4 | 7 |
| Clitellata | Oligochaeta | 288 | 312 | 261 | 861 |
| Diplopoda | Eugnatha | 3 | 21 | 3 | 27 |
| Gastropoda | Gastropoda | 14 | 30 | 62 | 106 |
| Insecta | *Acanthaspis* sp. | 5 | 4 | 4 | 13 |
|  | Blattoidae | 38 | 27 | 91 | 156 |
|  | Dermaptera & Orthoptera | 447 | 208 | 115 | 770 |
|  | Erotylidae | 3 | 18 | 15 | 36 |
|  | Formicidae | 197 | 274 | 303 | 774 |
|  | *Gonocephalum* sp. | 593 | 143 | 233 | 969 |
|  | Lepidoptera | 7 | 18 | 10 | 35 |
|  | Scarabaeidae | 21 | 16 | 71 | 108 |
|  | Termitoidae | 6 | 17 | 43 | 66 |
|  | Zygentoma | 7 | 1 | 2 | 10 |
| Malacostraca | Brachyura | 15 | 3 | 0 | 18 |
|  | Isopoda | 99 | 57 | 297 | 453 |
| Mammalia | *Mus* sp. | 0 | 0 | 1 | 1 |
| Reptilia | *Bungarus caeruleus* | 0 | 0 | 1 | 1 |
|  | *Echis carinatus* | 19 | 3 | 0 | 22 |
|  | *Eutropis* spp. | 4 | 2 | 3 | 9 |
|  | *Hemidactylus albofasciatus* | 23 | 1 | 0 | 24 |
|  | *Hemidactylus* sp. 1 | 2 | 0 | 0 | 2 |
|  | *Ophisops* spp. | 1 | 1 | 0 | 2 |
|  | Typhlopidae | 0 | 0 | 1 | 1 |
| **Total** | | **2405** | **1421** | **1912** | **5738** |

**Appendix S3 (a).** The effect of land-use change and rock size on the encounter rate of rocks from the results of negative binomial GLM. Rows highlighted in bold indicate coefficients whose 95% CI do not overlap zero and thus were inferred to significantly influence the response variable.

| **Term** | **Coefficient** | **95% CI** | ***p*** |
| --- | --- | --- | --- |
| **Intercept: (Plateau & Small Size)** | **3.9** | **3.63 – 4.2** | **< 0.001** |
| **Orchard** | **0.42** | **0.01 – 0.85** | **0.047** |
| **Paddy** | **-1.82** | **-2.29 – -1.35** | **< 0.001** |
| **Medium Size** | **0.87** | **0.47 – 1.26** | **< 0.001** |
| Large Size | -0.06 | -0.46 – 0.35 | 0.78 |
| Orchard x Medium Size | -0.26 | -0.84 – 0.33 | 0.393 |
| **Paddy x Medium Size** | **1.24** | **0.61 – 1.86** | **< 0.001** |
| Orchard x Large Size | 0.02 | -0.58 – 0.61 | 0.956 |
| **Paddy x Large Size** | **2.59** | **1.96 – 3.22** | **< 0.001** |

**S3 (b).** Estimated marginal means for factor combinations in negative binomial GLM assessing the effect of land-use change and rock size on the encounter rate of rocks. Rows highlighted in bold indicate estimated marginal means whose 95% CI do not overlap zero and thus were inferred to significantly influence the response variable.

| **Size** | **Contrast** | **Estimate** | **SE** | ***z*** | ***p*** |
| --- | --- | --- | --- | --- | --- |
| Small | Plateau – Orchard | -0.42 | 0.21 | -1.99 | 0.12 |
|  | **Plateau – Paddy** | **1.82** | **0.24** | **7.65** | **< 0.001** |
|  | **Orchard – Paddy** | **2.25** | **0.25** | **9.16** | **< 0.001** |
| Medium | Plateau – Orchard | -0.17 | 0.21 | -0.81 | 0.7 |
|  | **Plateau – Paddy** | **0.58** | **0.21** | **2.76** | **0.02** |
|  | **Orchard – Paddy** | **0.75** | **0.22** | **3.42** | **< 0.01** |
| Large | Plateau – Orchard | -0.44 | 0.21 | -2.06 | 0.0976 |
|  | **Plateau – Paddy** | **-0.77** | **0.21** | **-3.62** | **< 0.001** |
|  | Orchard – Paddy | -0.33 | 0.22 | -1.49 | 0.3 |

**Appendix S4.** The effect of land-use change on the proportion of detections of the focal species from the results of binomial GLMs (with a mixed bias-reduction adjusted scores approach for the models of *H. albofasciatus* and *E. carinatus*). Rows highlighted in bold indicate coefficients whose 95% CI do not overlap zero and thus were inferred to significantly influence the response variable.

| **Species** | **Land use type** | **Coefficient** | **95% CI** | ***p*** | ***Nagelkerke’s R^2^*** |
| --- | --- | --- | --- | --- | --- |
| *G. seshachari* | **Intercept: Plateau** | **-7.16** | **-8.95 – -6.03** | **<0.001** | **0.33** |
|  | Orchard | 0.99 | -0.48 **–**2.91 | 0.23 |  |
|  | **Paddy** | **2.88** | **1.67 – 4.71** | **<0.001** |  |
| *H. albofasciatus* | **Intercept: Plateau** | **-4.73** | **-5.14 – -4.31** | **<0.001** | **0.58** |
|  | **Orchard** | **-2.83** | **-4.48 – -1.18** | **<0.001** |  |
|  | **Paddy** | **-3.43** | **-6.23 – -0.63** | **0.02** |  |
| *E. carinatus* | **Intercept: Plateau** | **-4.87** | **-5.32 – -4.43** | **<0.001** | **0.42** |
|  | **Orchard** | **-1.84** | **-2.98 – -0.7** | **<0.01** |  |
|  | **Paddy** | **-3.29** | **-6.09 – -0.48** | **0.02** |  |

**Appendix S5.** Figure showing the number of different taxa belonging to the classes represented in the boxes on the left, encountered under rocks of varying sizes across land-use types. The number of individuals is depicted except in the cases of ants (Formicidae), termites (Termitoidae) and mites (Acariformes); their presence-absence data was considered.

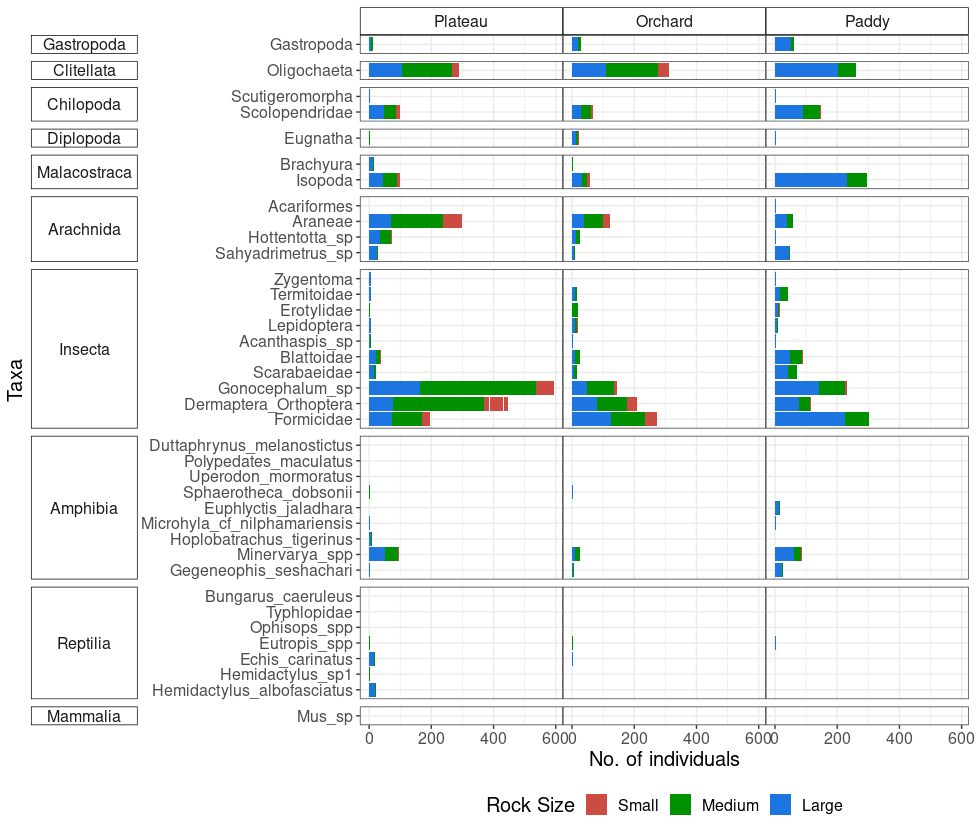

**Appendix S6.** Figure showing the rock dwelling animals composition across the land use types and rock size classes in the lateritic plateaus of the northern Western Ghats. Note that empty combinations are not shown in the figure. Vertical bars indicate the number of taxa recorded in each unique combination of land-use type and rock-size class shown with the connected points.

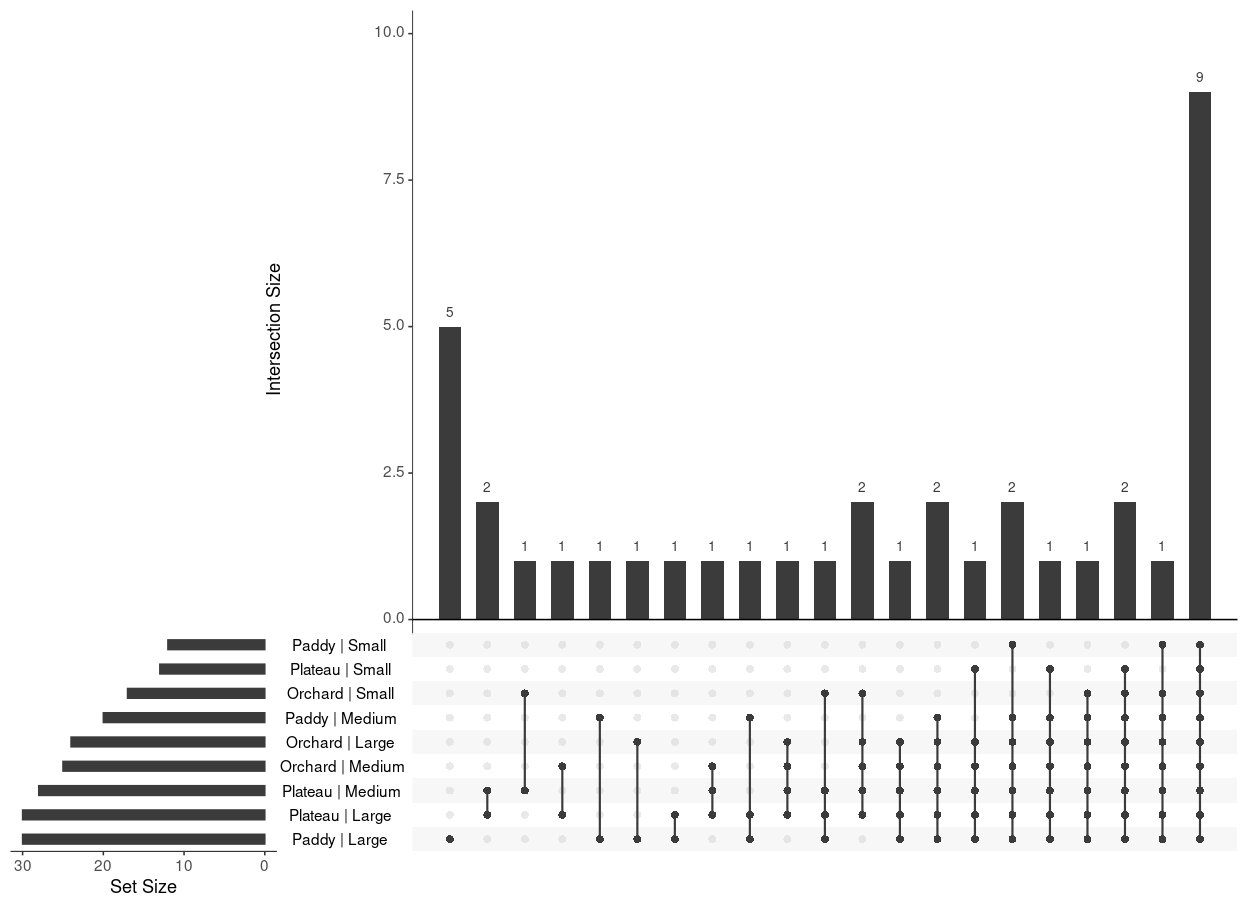

**Appendix S7.** Barplots showing the pairwise contrast coefficients from the generalized linear models used to assess the influence of land-use type on the encounter rate of animal groups with >100 individuals detected during the whole survey. Significance levels: ***p < .001, **p < .01, and *p < .05.

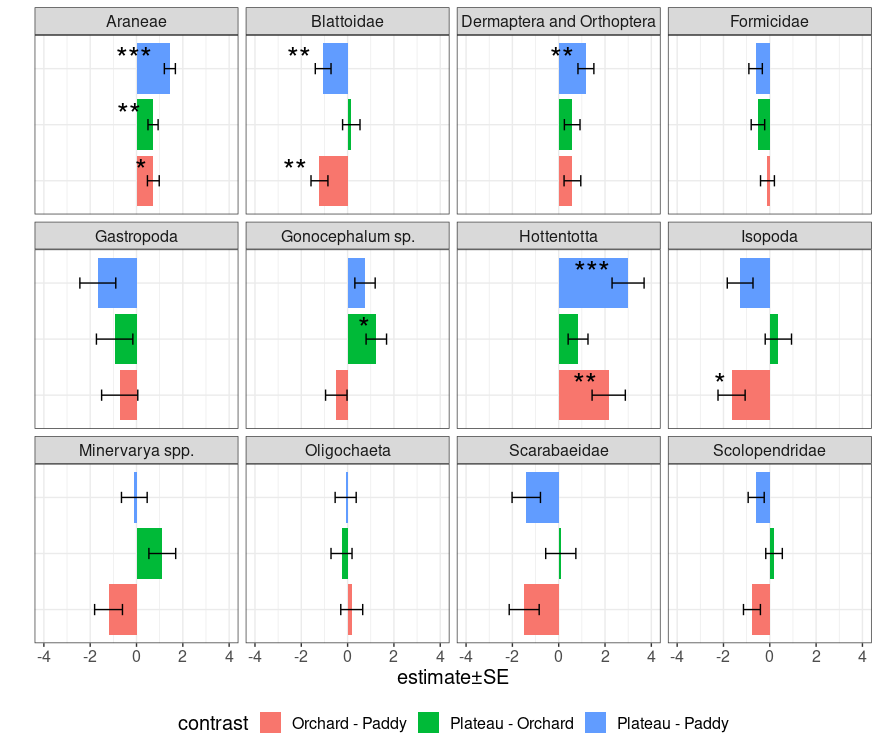

**Appendix S8.** Figure showing the use of rocks sourced from the plateau in paddy fields. Figure (a) shows an active paddy with rocks lined up to demarcate paddy boundaries; (b) and (c) shows gradual displacement of the boundary rocks creating scattered boulder grounds with wet soil after paddy abandonment. Photographs by V. Jithin.

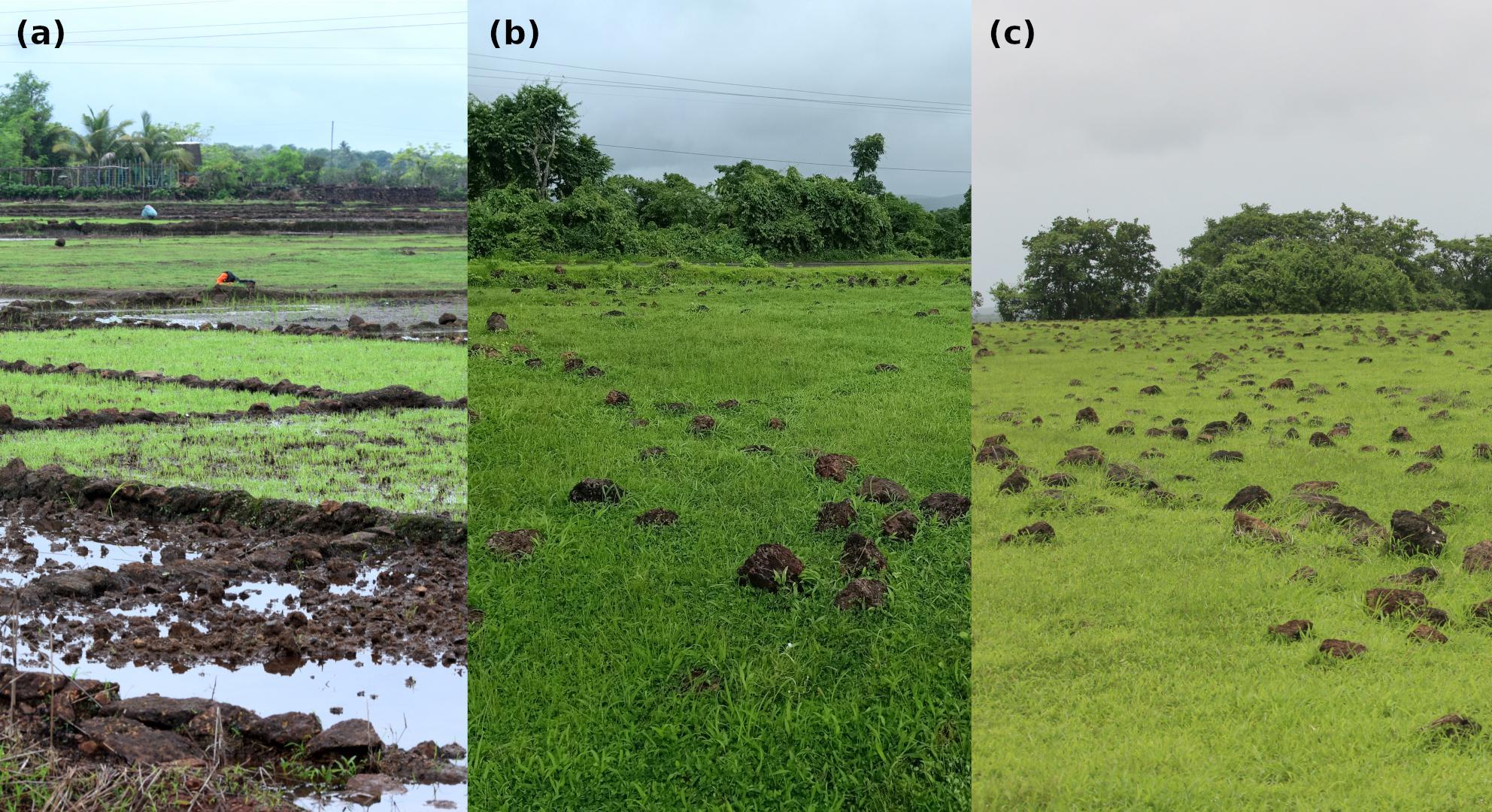

**Appendix S9.** Figure (a) showing a newly blasted plateau with plantation pits along a water diversion canal and (b) showing an establi37shed mango orchard with boundary rock wall around the property. Photographs by V. Jithin.

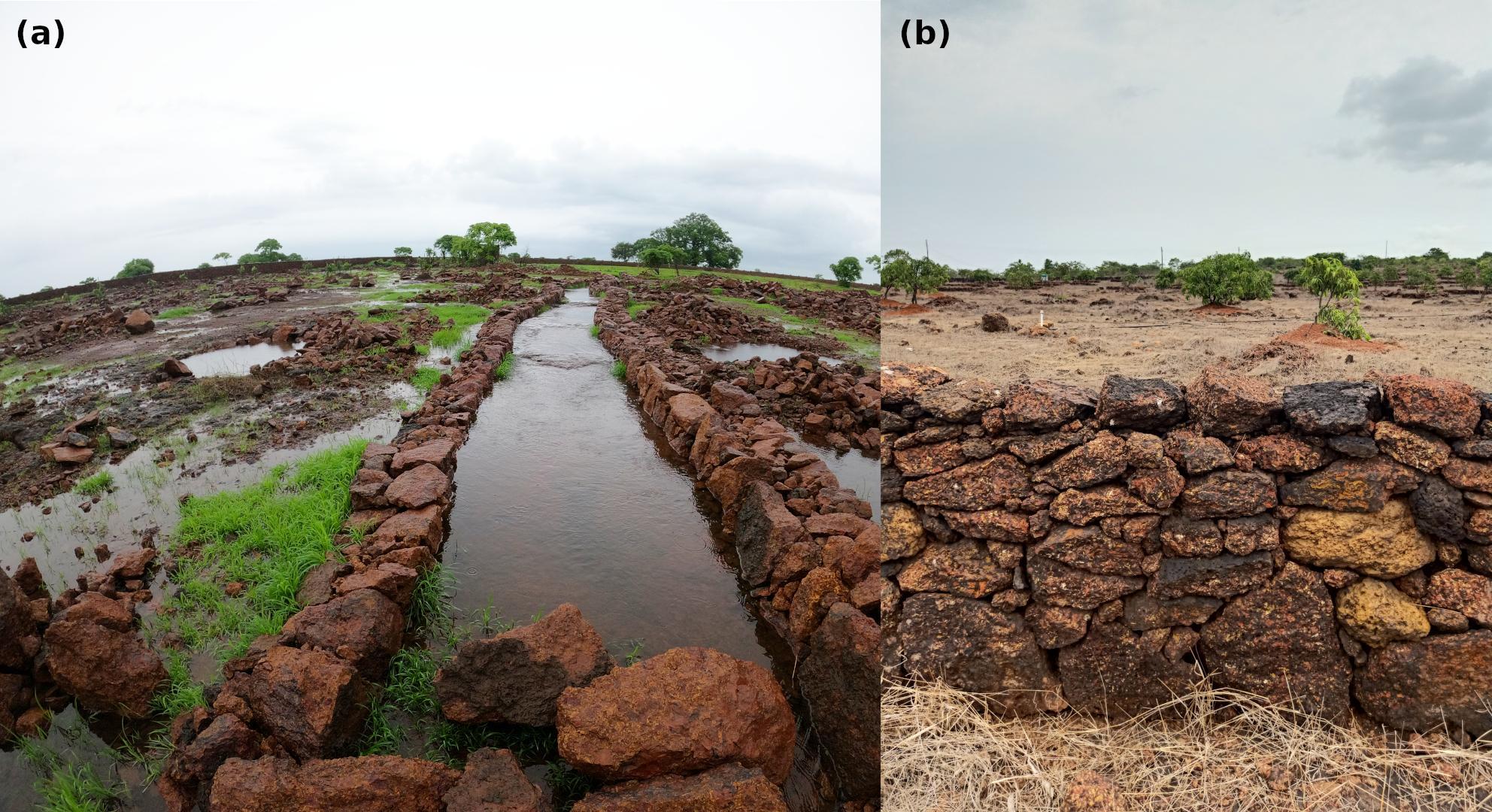
